# Leveraging Disease Association Degree for High-Accuracy MicroRNA Target Prediction

**DOI:** 10.1101/2025.11.06.686936

**Authors:** Baiming Chen

**Affiliations:** School of Medicine, The Chinese University of Hongkong, Shenzhen, Shenzhen, China

**Keywords:** microRNA, Language Model, Deep Learning

## Abstract

MicroRNAs (miRNAs) regulate gene expression by binding to mRNAs, inhibiting translation, or promoting mRNA degradation. Accurate identification of functional miRNA-target interactions (MTIs), typically validated by methods like western blot or reporter assay, remains challenging due to the scarcity of experimental data compared to the vast number of sequence-based predictions. This study pioneers a novel approach focusing solely on the disease association degree between miRNAs and their target genes. We propose that this single feature is sufficient for distinguishing experimentally validated functional MTIs from sequence-based predicted MTIs in a binary classification task. To quantify miRNA-gene disease association, we fine-tuned Sentence-BERT to generate disease description embeddings and compute their semantic similarity. Remarkably, using only disease association features, miRTarDS achieved an F1 score of 0.88 on the task of distinguishing functional from predicted MTIs in the external validation set. The approach also exhibits generalizability across different gene-disease association databases. This study demonstrates disease association as a powerful, independent dimension for prioritizing high-confidence functional MTIs.

## 1 Introduction

In 1993, miRNA (Lin-4) was first discovered and found to be critical for the timing of embryonic development in C. elegans [Lee et al., 1993]. Since then, a large number of miRNAs have been discovered with the development of high-throughput bioinformatics technologies [Van Kouwenhove et al., 2011]. miRNAs are classified as small non-coding RNAs (ncRNAs), and play a crucial role in the regulation of various physiological processes, such as cell growth, development, differentiation, and apoptosis [Cheng et al., 2005], as well as pathological processes [Hill and Tran, 2021]. Furthermore, miRNAs are promising biomarker candidates for various diseases [Pritchard et al., 2012]. Therefore, the identification of microRNA Target Interactions (MTIs) holds biological significance. Recently, the agoTRIBE has enabled single-cell resolution profiling of MTIs, overcoming the limitations of bulk sequencing, allowing scientists to gain deeper insights into miRNA functions across different cell types and states [Sekar et al., 2024].

There are many existing MTI prediction databases. miRWalk identifies consecutive complementary subsequences between miRNAs and genes to construct a database for MTIs prediction in different species [Sticht et al., 2018]. miRDB considers miRNA overexpression and CLIP-seq data, utilizing a support vector machine (SVM) for MTI prediction [Chen and Wang, 2020]. TargetScan predicts MTIs based on seed match and conservativeness analysis in different mammalians [Agarwal et al., 2015]. StarBase explores miRNA-mRNA and miRNA-lncRNA interaction networks based on CLIP-Seq data [Li et al., 2014]. The RNA22 is designed for the identification of miRNA binding sites and their corresponding miRNA and mRNA complexes [Miranda et al., 2006].

Several well-established MTI prediction tools exist, such as the classic tool miRanda [Enright et al., 2003] and PITA [Kertesz et al., 2007]. miRanda predicts MTIs by assessing sequence complementarity between miRNA and mRNA as well as the free energy of the resulting duplex structure. PITA predicts miRNA targets by considering both the binding free energy and the accessibility of the target site within the mRNA secondary structure. Recent advances have also introduced meachine learning method for prediction. miRAW employs stacked autoencoders to learn features directly from binary-encoded miRNA-mRNA sequences and relies on site accessibility filtering [Pla et al., 2018]. TargetNet innovates through alignment-enhanced encoding of extended seed regions and a ResNet architecture with 1D convolutions, enabling explicit modeling of binding patterns and quantitative distinction of high-functional targets [Min et al., 2022].

Although existing tools effectively leverage sequence features, they may overlook the functional information beyond sequence features. The previous research indicates that the dysregulation of miRNA and target genes often jointly drive specific disease phenotypes [Kloosterman and Plasterk, 2006]. For example, Rassenti et al. found that loss of miR-15/16 promotes overexpression of BCL2 and ROR1, each of them contributes to leukemia pathogenesis [Rassenti et al., 2017]. Since miRNAs and their target genes co-participate in disease pathways, MTIs’ disease association may be a potential feature for classifying functional and non-functional MTIs.

In recent years, miRNA-disease association (MDA) prediction [Huang et al., 2022] and disease gene prediction have advanced rapidly [Ata et al., 2021]. A notable example occurred in 2020, Graph Auto-Encoder for predicting miRNA-Disease Associations [Li et al., 2021] identified miR-122 as a top oesophageal cancer candidate. The following year, miR-122 was experimentally confirmed to inhibit KIF22, suppressing cancer cell progression [Wang et al., 2021]. Additionally, research on the association between miRNAs, genes, and related diseases has accumulated significantly. Consequently, the volume of data in databases that compile miRNA-disease associations and gene-disease associations from literature has become increasingly extensive. Examples include HMDD [Cui et al., 2024] and DisGeNE [Hu et al., 2025], which specifically curate miRNA-disease associations and gene-disease associations, respectively.

To quantify the disease association patterns between miRNAs and their target genes, we applied Sentence-BERT (SBERT) to compute semantic similarity between miRNA- and gene-associated diseases (Figure 1). SBERT improves the pre-trained BERT model by incorporating Siamese and triplet network structures [Reimers and Gurevych, 2019]. BERT, a bidirectional transformer model, captures the semantic meaning of each word based on its preceding and following context. Traditional BERT requires sentence pairs to be concatenated and processed together, making tasks like sentence similarity calculations computationally expensive. SBERT, by contrast, generates independent sentence embeddings that can be precomputed, allowing for sentence comparisons through simple similarity calculations such as cosine similarity. This approach significantly enhances inference speed. SBERT outperforms both BERT [Devlin et al., 2018] and RoBERTa [Liu et al., 2019] in terms of speed while maintaining comparable accuracy.

**Figure 1:**
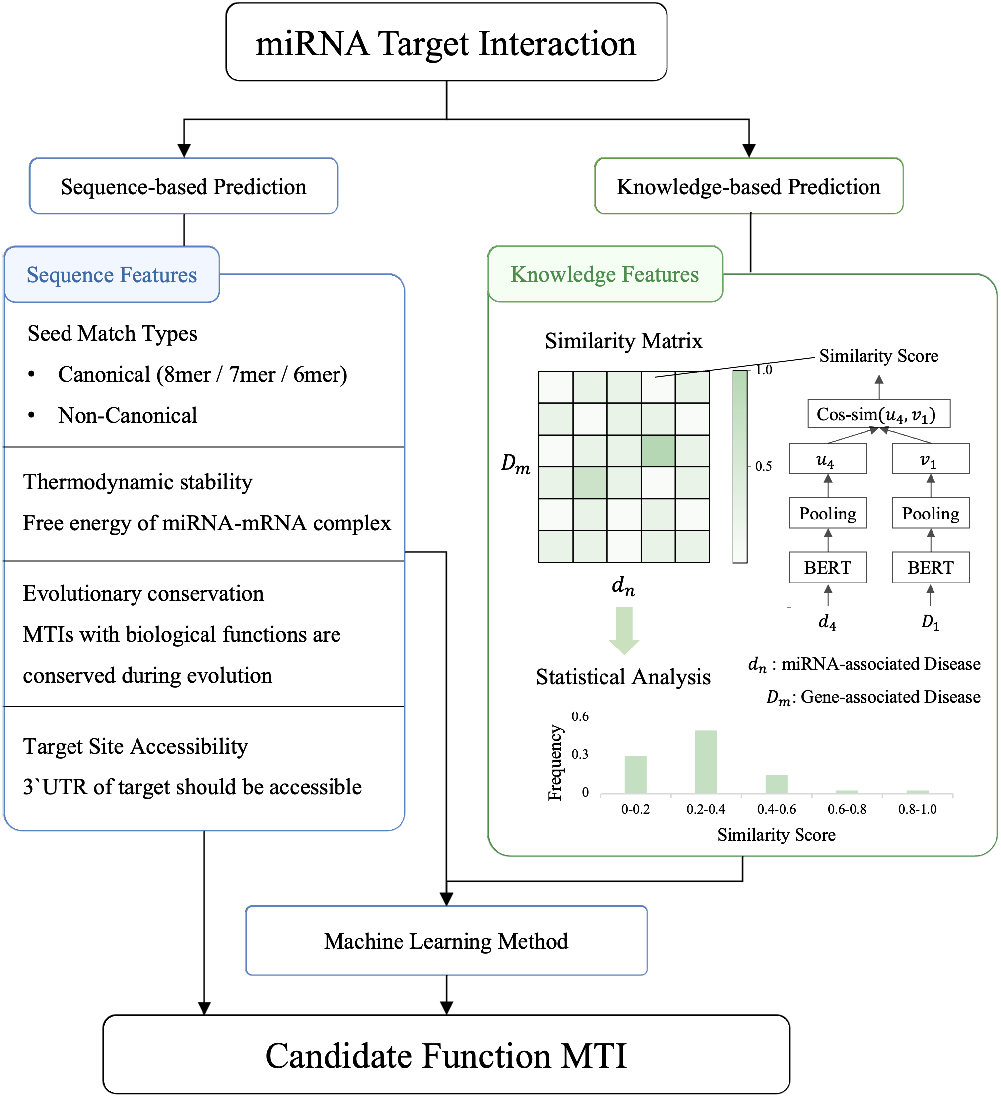
This figure illustrates the workflow of our research. Compared to the existing sequence-based prediction method (left), our method (right) proposes a novel method that directly utilizes the disease string as input. Here *d* and *D* denote sets of miRNA-associated and gene-associated disease descriptors, with *d*_*n*_ ∈ *d* and *D*_*m*_ ∈ *D* being individual descriptors. Embedding vectors are computed as *u*_*n*_ = *f*_SBERT_(*d*_*n*_) and *v*_*m*_ = *f*_SBERT_(*D*_*m*_).

Our central hypothesis is that functional MTIs exhibit distinct disease association patterns compared to sequence-based predictions. Therefore we applied a machine learning method that used the disease association between miRNA and gene as a feature, achieving an F1 score of 0.88 in a binary classification task. The experimental results confirm that functional MTI exhibits different disease association patterns from predicted MTIs (Supplementary Figure 1).

In summary, this study proposed and validated a novel method for quantifying miRNA-gene disease association using language model, demonstrating the utility of knowledge feature for enhanced MTI classification.

## 2 Materials and methods

### 2.1 Data Collection and Preprocessing

To generate a fine-tuning dataset for the SBERT model, we downloaded the MeSH (Medical Subject Headings) descriptors from the National Library of Medicine (https://meshb.nlm.nih.gov/). The MeSH descriptors are standardized terms utilized for indexing and retrieving biomedical and health-related information, forming a hierarchical vocabulary tree (Figure 3). To test the SBERT model performance, we obtained the BIOSSES dataset (Figure 2), which consists of biomedical statements with manually annotated text similarity scores provided by experts [Soğancioğlu et al., 2017]. It can be used to evaluate the biomedical text similarity calculation model.

**Figure 2:**
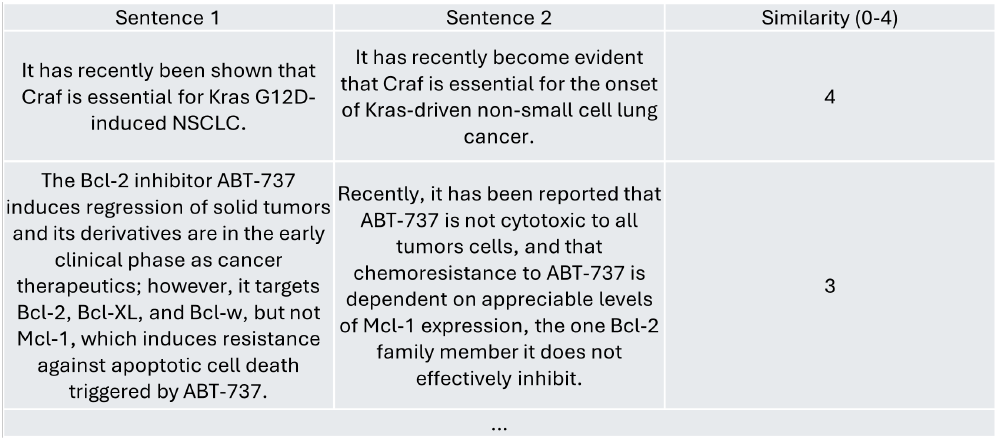
Example of BIOSSES dataset. BIOSSES dataset is a benchmark dataset specifically designed for evaluating semantic similarity calculation model in biomedical texts. The dataset consists of 100 sentence pairs extracted from biomedical literature, with each pair’s average semantic similarity score independently annotated by five experts.

**Figure 3:**
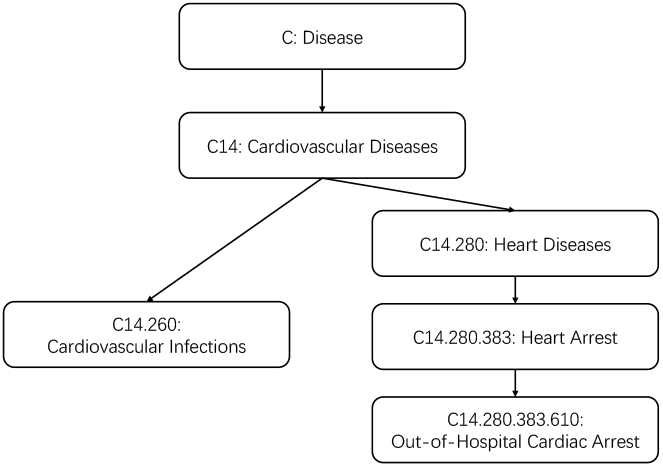
Disease descriptor ‘Cardiovascular Infections’ and ‘Out-of-Hospital Cardiac Arrest’ hierarchical structure in MeSH tree.

The successful application of machine learning classification techniques hinges on the availability of a sufficiently diverse and representative dataset, which is critical for ensuring that the trained model generalizes effectively to unseen data. To this end, we integrated data from multiple source databases. To obtain human functional MTIs and predicted MTIs, we collected miRTarBase [Cui et al., 2025], TarBase [Skoufos et al., 2024], TargetScan Agarwal et al. [2015], miRDB [Chen and Wang, 2020], and miRWalk [Sticht et al., 2018] (Table 1). miRTarBase and TarBase provide experimentally validated MTIs, TargetScan, miRDB and miRWalk provide predicted MTIs. For acquiring information on miRNA-associated and gene-associated diseases, we employed HMDD [Cui et al., 2024], DisGeNET [Hu et al., 2025] and KEGG [Kanehisa and Goto, 2000] databases. In the HMDD database, miRNA identifiers are provided without the ‘-3p’ or ‘-5p’ suffixes. Therefore, to maintain consistency across sources, we use pre-miRNA to replace mature miRNA. For instance, since hsa-miR-124-3p and TP53 are documented as a functional MTI in miRTarBase, the pair hsa-mir-124 and TP53 was classified as a functional MTI in our integrated dataset.

**Table 1:** Summary of MTIs and Source Databases Employed in the Study.

| Database | Human MTIs Count | Evidence Type | Source |
| --- | --- | --- | --- |
| miRTarBase 2025 | 8,168 | Experimental | https://mirtarbase.cuhk.edu.cn |
| TarBase 9.0 | 2,486 | Experimental | https://dianalab.e-ce.uth.gr/tarbasev9 |
| TargetScan 8.0 | 146,677 | Predicted | https://www.targetscan.org/vert_80 |
| miRDB 6.0 | 723,079 | Predicted | https://mirdb.org |
| miRWalk 3.0 | 8,073,040 | Predicted | http://mirwalk.umm.uni-heidelberg.de |

Our objective is to classify functional MTIs from predicted MTIs by leveraging disease association as a feature. Therefore, to construct a robust binary classification framework, we curated both positive and negative datasets based on the presence of experimental evidence. The positive set includes only high-confidence functional MTIs, while the negative set consists of computationally predicted MTIs that lack experimental support.

#### 2.1.1 Positive Dataset

The positive dataset comprises functional MTIs that have been experimentally validated through either reporter assays or western blot analyses. Reporter assays test the binding of a miRNA to the 3’UTR region of its target mRNA, while western blot confirms the resulting reduction in the expression level of the target protein. Both methods provide direct evidence of a specific interaction between a miRNA and its target gene.

#### 2.1.2 Negative Dataset

Although experimental techniques such as reporter assays and western blots are reliable for validating MTIs, they are resource-intensive, time-consuming, and unable to comprehensively screen all possible miRNA–mRNA pairs. Consequently, computational prediction serves as an essential tool for generating hypothetical MTIs on a large scale. These prediction tools analyze miRNA and mRNA sequences by incorporating factors such as seed sequence matching, target site accessibility, and binding free energy, etc., to score potential MTIs. The negative dataset consists of such predicted MTIs, which represent putative interactions that lack experimental confirmation.

#### 2.1.3 Training, Testing, and External Validation Splitting

To ensure a robust evaluation of model performance, we constructed a balanced dataset by combining all experimentally validated MTIs (positive set) with an equal number of randomly selected predicted MTIs (negative set), using a random seed of 42 for reproducibility. The combined MTIs were then split into an external validation set (10% of the total) and a remaining set (90%). The remaining set was further partitioned using 10-fold cross-validation for model training and internal evaluation. This stratified splitting strategy helps mitigate potential biases and enhances the generalizability of our results.

### 2.2 Fine-tuning Sentence-Bert Model

#### 2.2.1 Selection of Pretrained Model

The ideal sentence-bert model we seek must fulfill two primary criteria: exceptional performance in semantic similarity calculation tasks and rapid execution speed to facilitate large-scale data processing. Moreover, scalability for handling lengthy biomedical semantic strings is favorable. Based on the SBERT website (https://www.sbert.net/), we selected the ‘multi-qa-MiniLM-L6-cos-v1’ model for further investigation. This model excels in execution speed, and robust semantic similarity calculations, and supports extended input sequences.

#### 2.2.2 Synthetic Data for Fine-Tuning SBERT Model

The absence of SBERT models specialized for disease semantic similarity measurement, coupled with the scarcity of expert-annotated disease pairs, necessitates the creation of synthetic training data. Inspired by Wang et al. [2010], we generated disease similarity pairs using MeSH category C (diseases descriptors) to create a dataset containing pairs of diseases and their disease association degree, which calculated with the Lowest Common Ancestor (LCA) method. The distribution of disease association degree in the dataset we created closely mirrors the disease semantic similarity distribution produced by the ‘multi-qa-MiniLM-L6-cos-v1’ model without fine-tuning. Both types of distributions follow a log-normal distribution pattern. The data generation method is as follows:

- **Tree Number Representation:** For a MeSH descriptor, let 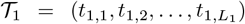 and 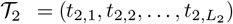 denote the hierarchical components of their tree numbers, where *L*_1_ and *L*_2_ are the depths of the two descriptors, respectively.
- Determine the depth of disease descriptors:

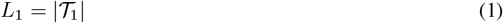

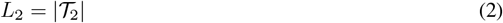

**Note:** |*T*_2_| stands for the number of hierarchical levels.
- Determine the depth of the lowest common ancestor (LCA):

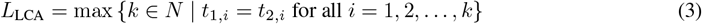

**Note:** *L*_LCA_ = 0 if no common ancestor exists.
- Calculate the total path depth:

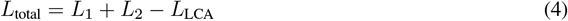
- Calculate the raw disease similarity:

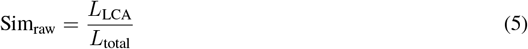

Calculate the adjusted disease similarity (decay factor *δ* = 0.8):

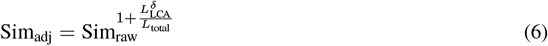
- **Example Calculation:** Consider Cardiovascular Infections (*T*_1_ = (C14, 260)) and Out-of-Hospital Cardiac Arrest (*T*_2_ = (C14, 280, 383, 610)) from the MeSH tree (Figure 3):

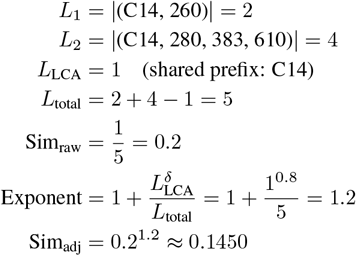

According to the calculated result, the weighted disease similarity between Cardiovascular Infections (C14.260) and Out-of-Hospital Cardiac Arrest (C14.280.383.610) is 0.1450. This similarity metric was further adjusted to better align with the distribution of the output generated by the ‘multi-qa-MiniLM-L6-cos-v1’ model, resulting in a dataset named ‘MeSHDS’. We utilized this dataset to fine-tune the SBERT model.

#### 2.2.3 Fine-Tuning Pretrained SBERT Model with MeSHDS

To enable the pre-trained SBERT model to effectively learn disease hierarchical relationships within the MeSH tree structure while preserving its intrinsic semantic understanding capabilities and mitigating catastrophic forgetting [Sun et al., 2019], we employed a carefully designed progressive fine-tuning strategy. This strategy utilizes a sequence of very small and monotonically decreasing learning rates, initialized at 5 × 10^−6^ and stepped down to 3 × 10^−6^, then 1 × 10^−6^.

Across all fine-tuning experiments, we maintained strict consistency in core hyperparameters and training configurations except for the specific learning rate schedule (i.e., the number of epochs allocated to each learning rate phase). This controlled approach isolates the effect of the learning rate progression. Hardware constraints permitting, we maximized the batch size per device to enhance gradient estimation stability during training. Optimization was performed using AdamW [Loshchilov and Hutter, 2017], known for its effective weight decay handling which aids in regularization. The optimization objective was the MSE loss between predicted and ground-truth disease similarity scores from MeSHDS.

For the *k*-th disease pair 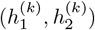:

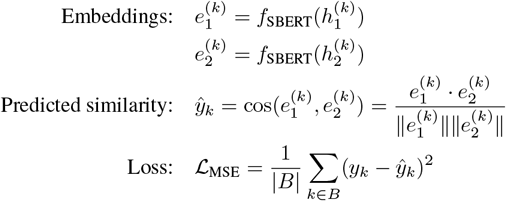

where 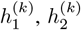 denote standardized disease terms from MeSH vocabulary (e.g., “Liver Neoplasms”, “Carcinoma, Hepatocellular”) in pair *k*. 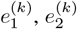 denote their SBERT embeddings. *y*_*k*_ ∈ [0, 1] is the ground-truth similarity. *B* represents training batch, and |*B*| represents the batch size.

The following learning rate schedules were evaluated:

LR 1: 5 × 10^−6^ for 1 epoch.

LR 2: 5 × 10^−6^ for 2 epochs.

LR 3: 5 × 10^−6^ for 2 epochs → 3 × 10^−6^ for 1 epoch.

LR 4: 5 × 10^−6^ for 2 epochs → 3 × 10^−6^ for 2 epochs.

LR 5: 5 × 10^−6^ for 2 epochs → 3 × 10^−6^ for 3 epochs.

LR 6: 5 × 10^−6^ for 2 epochs → 3 × 10^−6^ for 3 epochs → 1 × 10^−6^ for 1 epoch.

LR 7: 5 × 10^−6^ for 2 epochs → 3 × 10^−6^ for 3 epochs → 1 × 10^−6^ for 2 epochs.

### 2.3 Measurement of MTI Disease Association

To assess the phenotypic relatedness between miRNAs and genes based on their associated diseases, we quantified the semantic similarity between the sets of diseases linked to miRNAs and to genes. Using Sentence-BERT (SBERT), we generated embeddings for standardized disease descriptions and computed a pairwise cosine similarity matrix between all diseases in the miRNA-associated disease list and all diseases in the gene-associated disease list. The distribution of values within this similarity matrix was then analyzed to derive features characterizing the overall semantic relatedness between the disease profiles of the miRNA and gene.

#### Semantic String Embedding

For a miRNA-associated disease descriptor set *d* = {*d*_1_, *d*_2_, …, *d*_*n*_} and gene-associated disease descriptor set *D* = {*D*_1_, *D*_2_, …, *D*_*m*_}, we computed their embedding matrices:

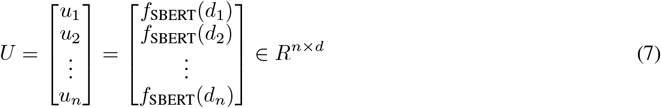

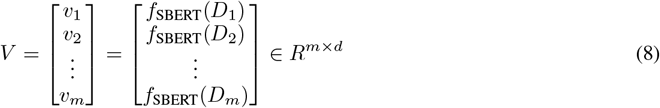

where *u*_*n*_, *v*_*m*_ ∈ *R*^*d*^ are *d*-dimensional embedding vectors produced by SBERT, and *d* = 384 in our implementation.

#### Similarity Matrix Calculation

The cosine similarity matrix between miRNA-associated disease and gene-associated disease *S* ∈ *R*^*n*×*m*^ is computed by:

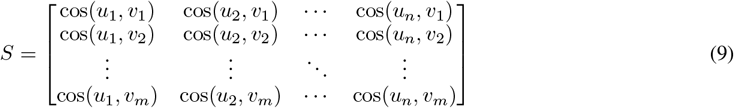

where the cosine similarity between two embeddings can be calculated by dot-product since ∥*u*_*n*_∥ = ∥*V*_*m*_∥ = 1:

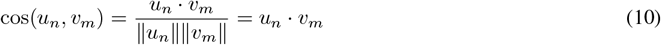

#### Feature Construction

The similarity matrix *S* is transformed into a normalized 5-dimensional feature vector *p*_sim_ representing the empirical distribution of similarity scores:

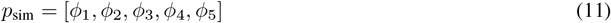

where

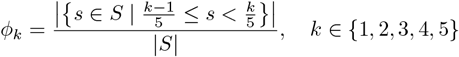

and |*S*| = *m* × *n* denotes the total number of elements in the matrix. *p*_sim_ forms a probability distribution with 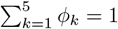.

### 2.4 Identifying Optimal Fine-tuned Models and Superior Classification Method

Functional MTIs and predicted MTIs were classified based on the disease association, with the classifier’s performance evaluated using the F1 score. The F1 score, calculated as the harmonic mean of precision and recall, was affected by different fine-tuned models and the choice of classification methods in this study. To ensure optimal performance, we conducted extensive tuning to identify the most suitable model and classification approach. We employed 7 machine learning algorithms, including Decision Tree, Gradient Boosting, Logistic Regression, Neural Network, Random Forest, Support Vector Machine (SVM), XGBoost to evaluate their performance through 10-fold cross-validation on the training set and further validating them on an external validation set.

## 3 Result

### 3.1 Disease Similarity Calculation Performance of Fine-Tuned SBERT Model

The “MeSHDS” dataset comprises 2,249,975 disease pairs along with their corresponding similarities (Figure 4), a substantial volume of data that ensures the high quality of fine-tuning. To test the performance of the fine-tuned SBERT model, we calculated Pearson’s correlation coefficient between the model output scores and scores from BIOSSES and MeSHDS (Figure 5). The results demonstrated that the model retained its original capacity for biomedical semantic similarity computation while successfully learning new knowledge through fine-tuning.

**Figure 4:**
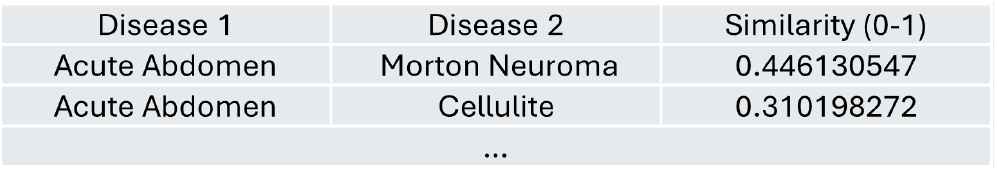
Example of MeSHDS dataset. Disease 1 and Disease 2 are derived from the MeSH Tree, and their disease association is computed using the LCA method described in section 2.2.2

**Figure 5:**
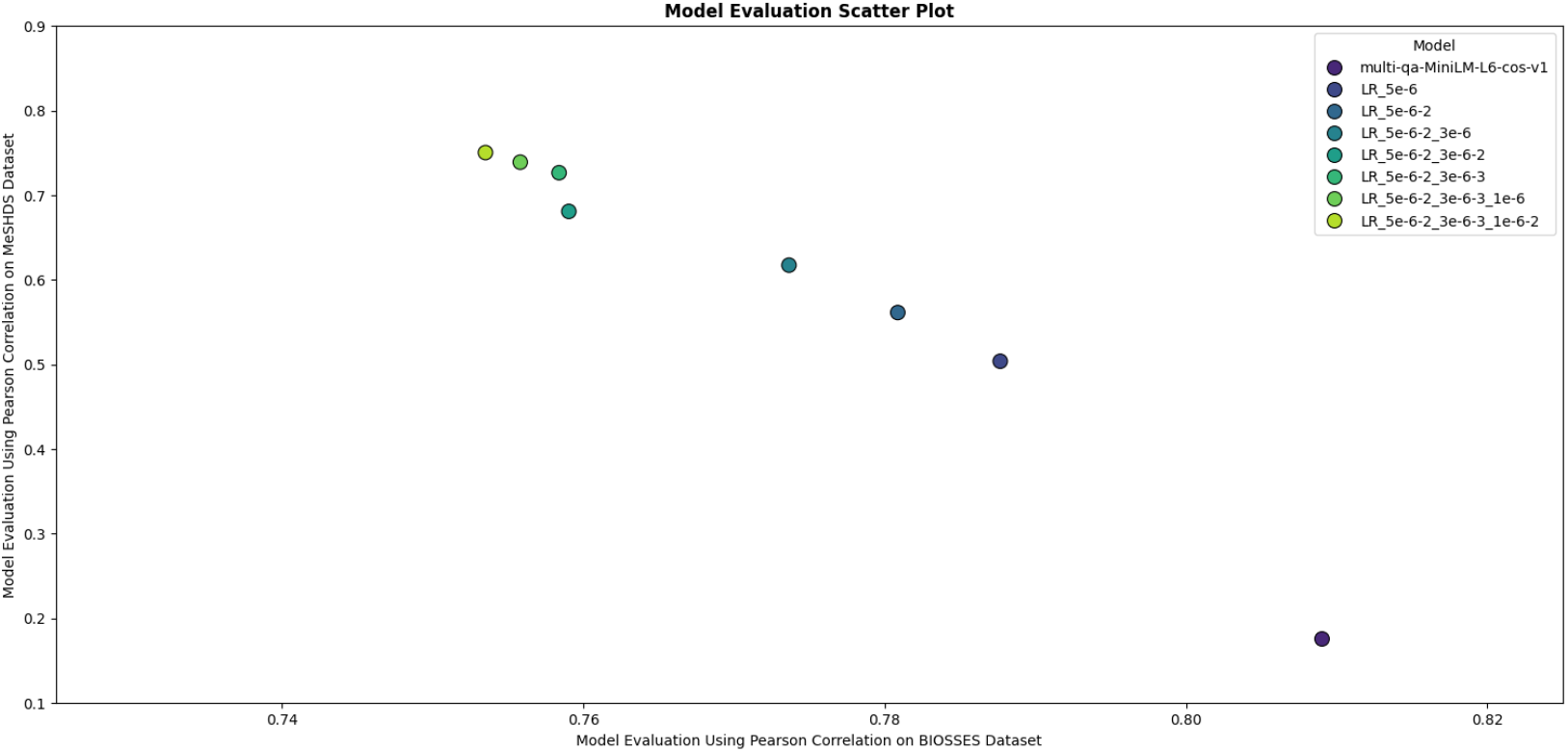
Model performance benchmark. The x-axis represents the correlation coefficient between the model-generated similarity scores and the BIOSSES dataset similarity scores, measuring the model’s ability to handle semantic similarity in the biomedical domain. The y-axis represents the correlation coefficient between the model-generated similarity scores and the MeSHDS dataset similarity scores, evaluating the fine-tuning effectiveness.

### 3.2 Fine-Tuned Model and Classification Method Selection

The F1 score was employed as the primary selection criterion to identify the optimal SBERT model and machine learning method for classifying functional MTIs and predicted MTIs. The pre-trained model fine-tuned using MeSHDS for two rounds, with a learning rate of ‘5e-6’, followed by an additional two rounds of fine-tuning at a learning rate of ‘3e-6’ demonstrated remarkable performance improvement, achieving an 5% increase compared to the pre-trained model. The utilization of Neural Network classification led to optimal overall performance, as evidenced by an external F1 Score of 0.88 (Table 2). After comparison, we selected the ‘LR 4’ (LR_5e_6-2_3e_6-2) model and employed the Neural Network for further investigation.

**Table 2:**
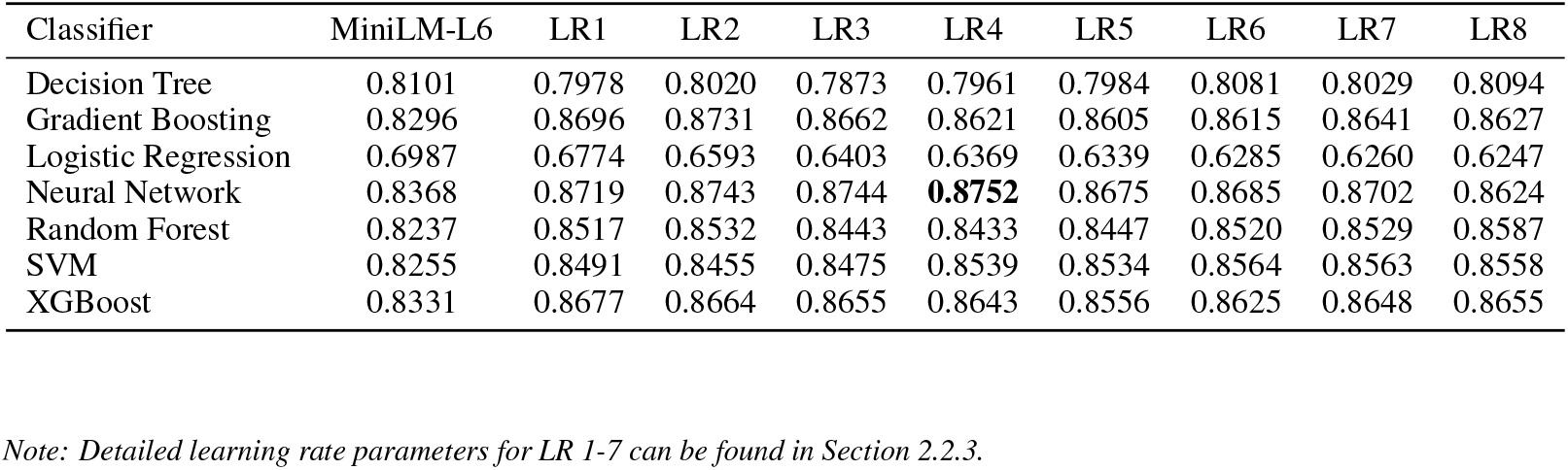
F1 Scores for Different SBERT Models and Machine Learning Classifiers.

| Classifier | MiniLM-L6 | LR1 | LR2 | LR3 | LR4 | LR5 | LR6 | LR7 | LR8 |
| --- | --- | --- | --- | --- | --- | --- | --- | --- | --- |
| Decision Tree | 0.8101 | 0.7978 | 0.8020 | 0.7873 | 0.7961 | 0.7984 | 0.8081 | 0.8029 | 0.8094 |
| Gradient Boosting | 0.8296 | 0.8696 | 0.8731 | 0.8662 | 0.8621 | 0.8605 | 0.8615 | 0.8641 | 0.8627 |
| Logistic Regression | 0.6987 | 0.6774 | 0.6593 | 0.6403 | 0.6369 | 0.6339 | 0.6285 | 0.6260 | 0.6247 |
| Neural Network | 0.8368 | 0.8719 | 0.8743 | 0.8744 | **0.8752** | 0.8675 | 0.8685 | 0.8702 | 0.8624 |
| Random Forest | 0.8237 | 0.8517 | 0.8532 | 0.8443 | 0.8433 | 0.8447 | 0.8520 | 0.8529 | 0.8587 |
| SVM | 0.8255 | 0.8491 | 0.8455 | 0.8475 | 0.8539 | 0.8534 | 0.8564 | 0.8563 | 0.8558 |
| XGBoost | 0.8331 | 0.8677 | 0.8664 | 0.8655 | 0.8643 | 0.8556 | 0.8625 | 0.8648 | 0.8655 |

Note: Detailed learning rate parameters for LR 1-7 can be found in Section 2.2.3.

### 3.3 Comparison with Other miRNA Target Predictors

To contextualized the performance of our knowledge-driven approach, miRTarDS, we compared it against several widely recognized and established sequence-based miRNA target prediction algorithms: miRanda [Miranda et al., 2006], TargetScan [Agarwal et al., 2015], PITA [Kertesz et al., 2007], miRDB [Chen and Wang, 2020], miRAW [Pla et al., 2018], and TargetNet [Min et al., 2022]. These methods represent significant contributions to the field of miRNA target prediction. Performance metrics for the sequence-based predictors were excerpted from the comprehensive evaluation conducted by Pla et al. [2018] and Min et al. [2022], where the optimal reported result for each algorithm was displayed. We also compared the performance of our method with that of the MISIM algorithm [Wang et al., 2010] in calculating disease associations.

The miRTarDS dataset tests the model’s ability to classify functional MTIs and computational predicted MTIs. miRAW tests the model’s ability to classify functional MTIs and non-functional MTIs. In miRAW, positive examples are built from experimentally validated miRNA–mRNA interactions (Diana TarBase / miRTarBase). Negative examples were drawn from experimentally verified non-target miRNA–mRNA pairs by scanning the 3’UTR with a 30-nt sliding window and keeping only regions that could form a stable duplex according to RNACoFold. It is important to note a potential sampling discrepancy in miRAW: while the dataset is approximately balanced at the sequence-site level, a significant imbalance exists at the level of miRNA–gene pairs, where positive samples substantially outnumber negative ones. This imbalance may introduce a slight bias when evaluating the performance of disease-association methods on this dataset. However, due to current limitations in data availability, there is a general lack of systematic experimental documentation for non-functional MTIs, making this issue difficult to resolve currently.

As shown in Table 3, miRTarDS demonstrated strong performance on both the miRAW dataset and miRTarBase dataset. These results underscore the efficacy of leveraging biological knowledge for the miRNA target interaction (MTI) prediction task.

**Table 3:** Performance Comparison of miRNA Target Prediction Methods.

| Method | Dataset | Accuracy | Precision | Recall | Specificity | F-score |
| --- | --- | --- | --- | --- | --- | --- |
| **Sequence-Feature Based Methods** |  |  |  |  |  |  |
| miRanda | miRAW | 0.5119 | 0.5205 | 0.3011 | 0.7226 | 0.3815 |
| Targetscan | miRAW | 0.5766 | 0.6077 | 0.4325 | 0.7208 | 0.5053 |
| PITA | miRAW | 0.5128 | 0.5461 | 0.1515 | 0.8741 | 0.2371 |
| miRDB | miRAW | 0.5493 | **0.7500** | 0.1478 | **0.9507** | 0.2470 |
| miRAW | miRAW | 0.7055 | 0.6749 | 0.7923 | 0.6186 | 0.7289 |
| TargetNet | miRAW | **0.7251** | 0.6572 | **0.9411** | 0.5091 | **0.7739** |
| **Disease-Association Feature Based Methods** |  |  |  |  |  |  |
| MISIM | miRAW | 0.6556 | 0.6674 | 0.6550 | 0.6700 | 0.6473 |
|  | miRTarDS | 0.7988 | 0.7989 | 0.7988 | 0.7990 | 0.7987 |
| miRTarDS | miRAW | 0.5844 | **0.9549** | 0.5844 | 0.4889 | 0.7168 |
|  | miRTarDS | **0.8753** | 0.8758 | **0.8753** | **0.8601** | **0.8752** |

Note: The miRTarDS dataset evaluates the model’s performance in distinguishing functional MTIs from computationally predicted ones, while miRAW tests its ability to differentiate between functional and non-functional MTIs.

## 4 Discussion

The present study introduces a method to refine sequence-based predicted miRNA-target interactions (MTIs) by incorporating the semantic similarity between diseases associated with miRNAs and diseases associated with target genes. This approach ultimately supports the development of a novel database for predicting MTIs with functional similarity and identifying detailed associated diseases.

Our analysis of disease similarity distributions (Supplementary Figure 1) revealed statistically significant differences between functional MTIs (validated by western blot or reporter assay from miRTarBase) and predicted MTIs (de novo predictions from miRWalk and miRDB). Specifically, functional MTIs exhibited a lower frequency (p=4.10e-47, independent samples t-test) in the low-similarity range (0-0.2) and a higher frequency (p=6.22e-71) in the moderate-similarity range (0.2-0.4) compared to predicted MTIs. This demonstrates that functional MTIs are associated with significantly higher miRNA-gene disease similarity scores than purely predicted MTIs, confirming our hypothesis that disease association patterns differ between these MTI types.

Interestingly, based on our correlation analysis, we observed a positive association between the number of disease associations for both miRNAs and genes and the prediction accuracy of our model. Specifically, we found a correlation coefficient of 0.271 between miRNA-disease association counts (above the first quartile threshold) and prediction correctness (TF), while the correlation was slightly stronger (0.312) for gene-disease association counts. For miRNA-disease associations, prediction accuracy increased substantially from 77.3% in the lowest quartile (1.00-70.00 associations) to approximately 94-97% across the remaining three quartiles. Similarly, for gene-disease associations, we observed a progressive improvement in prediction accuracy from 74.4% in the lowest quartile (1.0-68.8 entries) to 98.2% in the highest quartile (382.0-2724.0 entries). These findings indicate that the availability of extensive disease association data for both miRNAs and genes contributes to more accurate predictions. They also highlight a limitation of our study: the relatively lower prediction accuracy for MTIs involving miRNAs or genes with limited disease association data.

We also integrated both disease association features and sequence-based features to classify functional MTIs and predicted MTIs. The sequence feature was represented by scores obtained from miRDB. As shown in Table 4, the model utilizing both feature types demonstrated a slight improvement in performance compared to using the disease association feature and sequence feature alone. The comparison results prove that the disease association degree is a very important dimension.

**Table 4:** Performance Comparison of Using Different Feature.

| Model | Accuracy | Precision | Recall | Specificity | F-score |
| --- | --- | --- | --- | --- | --- |
| **Sequence Features** |  |  |  |  |  |
| Random Forest | 0.5550 | 0.5556 | 0.5551 | 0.5640 | 0.5541 |
| SVM | 0.6196 | 0.6270 | 0.6196 | 0.7364 | 0.6135 |
| Neural Network | 0.6223 | 0.6271 | 0.6223 | 0.7133 | 0.6184 |
| **Disease Association Feature** |  |  |  |  |  |
| Random Forest | 0.8173 | 0.8192 | 0.8173 | 0.8043 | 0.8170 |
| Gradient Boosting | 0.8404 | 0.8435 | 0.8404 | 0.8016 | 0.8401 |
| Neural Network | 0.8418 | 0.8448 | 0.8418 | 0.8044 | 0.8414 |
| **Sequence Features & Disease Association Feature** |  |  |  |  |  |
| Random Forest | 0.8261 | 0.8280 | 0.8261 | 0.8071 | 0.8259 |
| Gradient Boosting | 0.8404 | 0.8435 | 0.8404 | 0.7990 | 0.8401 |
| Neural Network | 0.8472 | 0.8497 | 0.8472 | 0.8193 | 0.8469 |

We further investigated potential biases arising from the use of different disease association databases. Gene–disease associations for TP53, PTEN, KRAS, MYC, TNF, IL6, IL10, and CRP were manually curated from the KEGG database. These genes were selected to generate disease association features for 338 MTIs using both DisGeNET and KEGG. Pearson’s correlation coefficients between the feature sets derived from the two databases were computed for each MTI. The average correlation coefficient was approximately 0.97, and the classifier performances were highly consistent. These results indicate that the proposed method is robust and generalizable across different databases. Nevertheless, since KEGG contains fewer gene–disease associations and does not allow complete database downloads, DisGeNET remains the more practical choice for large-scale analyses.

## 5 Future Work

The NLP-based approach for calculating disease semantic similarity, which was fine-tuned on our generated dataset, exhibited strong performance, confirming its utility in biomedical contexts. This method offers a scalable strategy for estimating functional association or disease association across diverse biomolecules, thereby supporting the exploration of potential interactions. In addition to enabling interaction prediction based on physical and chemical properties, this approach introduces a novel perspective through the quantification of disease association degrees, thus enhancing the overall analytical framework. For instance, it could be leveraged existing protein-disease association databases such as the Proteome-Phenome Atlas [Deng et al., 2025] and DisGeNET [Hu et al., 2025]—to analyze protein-protein interactions (PPIs).

Moving forward, we plan to apply our model to all collected MTIs and establish a publicly accessible online database. This resource will provide high-accuracy predictions of functional MTIs, along with detailed associations between MTIs and related diseases as identified by our method, to support further research within the scientific community.

## 6 Acknowledgement

The authors declare that there are no conflicts of interest. The methodology described above is the subject of a pending patent application.

## Supplementary Figures

**Figure 1:**
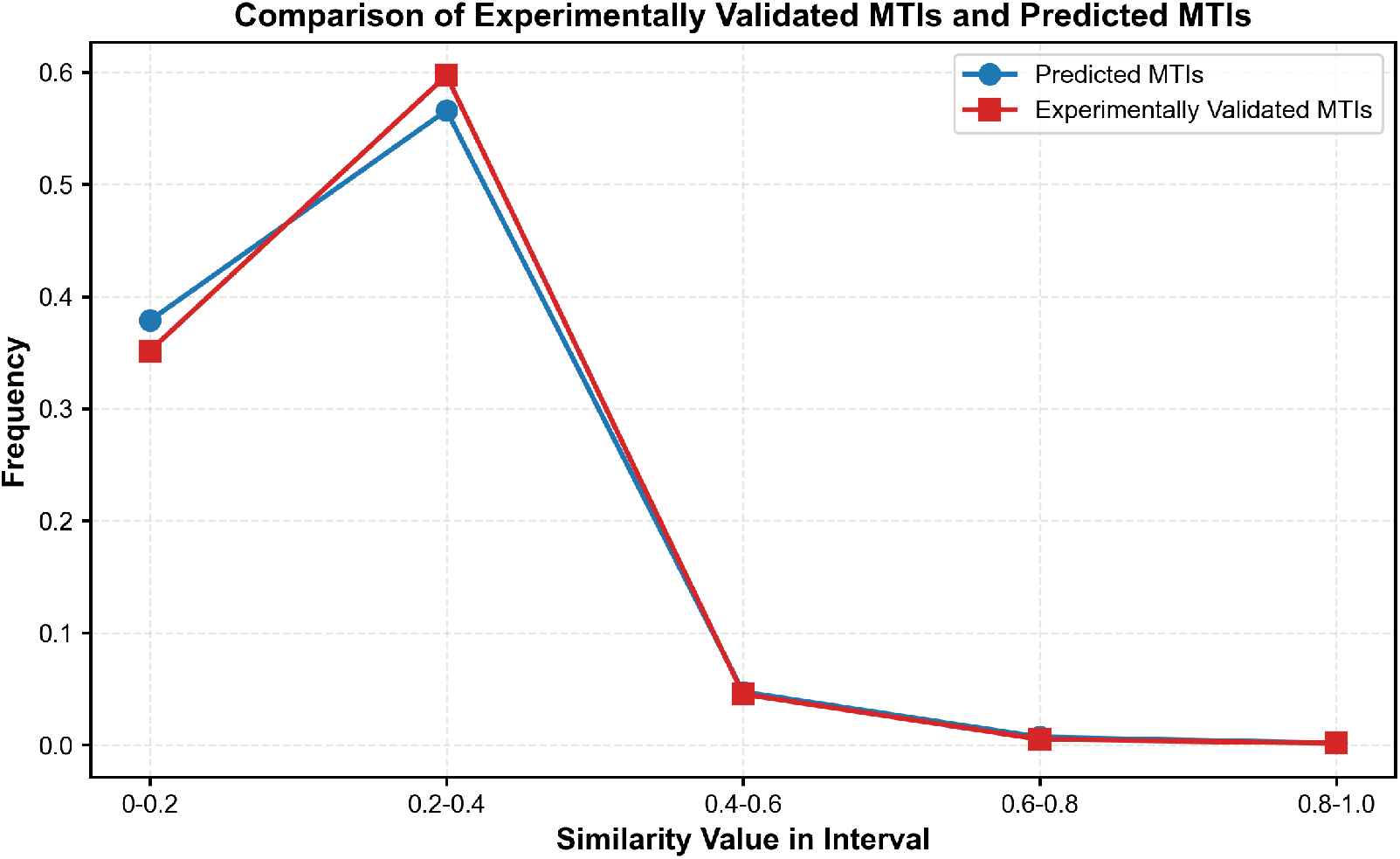
This figure presents the average frequency distribution of similarity values between miRNA-associated diseases and gene-associated diseases for both experimentally validated MTIs and predicted MTIs. Experimentally validated MTIs are confirmed using western blot or reporter assay, while predicted MTIs are derived from computational predictions based on sequence data. The x-axis represents the intervals of similarity values, and the y-axis indicates the average frequency of these similarity values within each interval. Specifically, functional MTIs exhibited a lower frequency (p=4.10e-47, independent samples t-test) in the low-similarity range (0-0.2) and a higher frequency (p=6.22e-71) in the moderate-similarity range (0.2-0.4) compared to predicted MTIs. The figure directly demonstrates that diseases associated with miRNAs and genes in experimentally validated MTIs show greater association compared to those in predicted MTIs.

